# ThermoTargetMiner as a proteome integral solubility alteration target database for prospective drugs against lung cancer

**DOI:** 10.1101/2024.08.06.606599

**Authors:** Hezheng Lyu, Hassan Gharibi, Bohdana Sokolova, Mücahit Varli, Anna Voiland, Brady Nilsson, Zhaowei Meng, Massimiliano Gaetani, Amir Ata Saei, Roman A. Zubarev

## Abstract

Knowledge of the targets of therapeutic compounds is vital for understanding their action mechanisms and side effects, but such valuable data is seldom available. The multiple complementary techniques needed for comprehensive target characterization must combine data reliability with sufficient analysis throughput. Here, we leveraged the Proteome Integral Solubility Alteration (PISA) assay to characterize the targets of 67 approved and experimental compounds against two main lung cancer subtypes, which is now provided through the website https://thermotargetminer.serve.scilifelab.se/app/thermotargetminer. Novel target candidates (pro-targets) were found for 77% of the tested molecules. Comparison of the protein solubility shifts in lysate vs. living cells highlighted the targets directly interacting with the compounds. We verified that the drug PEITC exerts cytotoxicity through the inhibition of a pro-target PAFAH1B. As PISA is now joining the arsenal of fast and reliable target characterization techniques, the presented database, ThermoTargetMiner, will become a useful resource in lung cancer research.

## Introduction

Lung cancer is one of the most prevalent cancer types, with approximately 2.2 million new cases every year resulting in 1.8 million deaths worldwide^1^. As the main lung cancer subtype, non-small-cell lung cancer (NSCLC) comprises 85% of all cases. NSCLC is divided into 3 types: squamous cell carcinoma, adenocarcinoma and large cell (undifferentiated) carcinoma^2^. Small-cell lung cancer (SCLC) accounts for the rest of lung cancer cases and exhibits neuroendocrine properties. Being strongly associated with tobacco exposure, it is highly aggressive and rapidly growing.

Small-molecule chemotherapy is the main approach to managing lung cancer in a stage-specific manner. Surgery can effectively manage tumor removal for the majority of early-stage NSCLC^3^. Neoadjuvant and adjuvant chemotherapy delivered before or after surgery is widely used in stage II and stage III NSCLC and is the primary approach for metastatic NSCLC. Recently, definite chemoradiotherapy combined with immune checkpoint inhibitors (ICIs) has become the preferred treatment for unresectable stage III NSCLC. For SCLC, localized cases are usually treated with surgery and concurrent chemoradiotherapy^4^. Adding ICIs to the conventional first-line platinum-based chemotherapy is the recommended approach for treating newly diagnosed metastatic SCLC.

The 5-year survival of NSCLC is around 60-70% for stage I, 40-50% for stage II, 5-25% for stage III and less than 1% for stage IV^2^. SCLC is also characterized by poor prognosis and has a remarkable tendency for early metastasis. Most patients respond to treatments only temporarily, which leads to a median survival of less than two years for those with early-stage SCLC and around one year for those with metastatic disease^4^.

Due to the limited survival of lung cancer patients, there is an urgent need for new drug development in this area. In April 2024, there were 317 registered small molecule drugs for lung cancer and 2795 registered clinical trials for such drugs on http://clinicaltyrials.gov. Over 70% of the clinical trials were in phase II and phase III. The primary reason for failure in both phase II and phase III is drug inefficacy ^5^. In many cases this failure is due to the poor knowledge of the drug action mechanism. The latter implies possessing solid information on drug targets, including the residence time of the drug on its target^6^. Another important reason for clinical trial failure is unacceptably high drug toxicity, resulting from the engagement of unintended off-targets by the drug or its metabolites^7,8^. Therefore, the pharmaceutical industry heavily invests in target characterization before clinical trials. To reduce the expenses and speed up drug development, the techniques for drug target characterization should combine the reliability of produced results with a reasonably high throughput. Another important desirable aspect of analysis is its proteome-wide nature, to reduce the risk of overlooking important (off-)targets.

In the last 10-15 years, several system-wide methods for dissecting drug targets have emerged. A well-known approach to probing drug-target interactions, thermal stability shift assay^9,10^, has been expanded to complex in-vitro and even in-vivo settings, while its first implementation, cellular thermal shift assay (CETSA^10^), required *a priori* knowledge of the target. The latter drawback was overcome in Thermal Proteome Profiling (TPP^11^ or MS-CETSA^10^) that employs mass spectrometry (MS) for a proteome-wide search of target candidates. These techniques detect the change in the melting temperature of a target protein upon binding to a small molecule. The main bottleneck of these MS-based approaches was their low throughput. In contrast, Proteome Integral Solubility Alteration (PISA) assay^12^, which analyzes the shift in protein solubility rather than that in melting temperature, offers at least an order of magnitude higher throughput. On top of that, PISA provides the possibility to employ different solubility modulators besides elevated temperature^12^, such as, e.g., organic solvents^13^ and kosmotropic salts^14^. When applied to a cell lysate, all the above approaches will highlight the proteins that bind directly to the drug, while the application to living cells also reveals the downstream proteins as well as the targets of drug metabolites^15^.

We have previously applied PISA for protein target identification and exploring mechanisms of small-molecule drugs^6,12^, biomarker discovery^16^, as well as for identification and prioritization of enzyme substrates^15,17^. In the current study, we tested the PISA performance as a higher-throughput technique for reliable drug target deconvolution on a larger set of drugs encompassing a diverse set of known drug targets, choosing lung cancer as the application area (**Figure 1**). For that purpose, we selected 67 therapeutic agents specifically designed or repurposed for lung cancer treatment and included both NSCLC and SCLC cell lines as disease models (A549 and NCI-H82 cells, respectively) to build an online expandable resource of drug targets called ThermoTargetMiner. Both the cell lysates and intact cells were treated by each drug and vehicle as control, PISA-processed, after which the proteomes were extracted and analyzed by a combination of liquid chromatography and tandem mass spectrometry (LC-MS/MS). The shifts of the protein PISA signals quantified using the tandem mass tag (TMT) were then analyzed. From the previous similar efforts with drug target identification by expression proteomics^18^, we expected that the datasets for the drugs with the same target would be found to be co-localized in hierarchical clustering, revealing similar drug mechanisms. However, in the PISA data drugs with the same target did not necessarily cluster together, which complicated the action mechanism determination.

**Figure 1.**
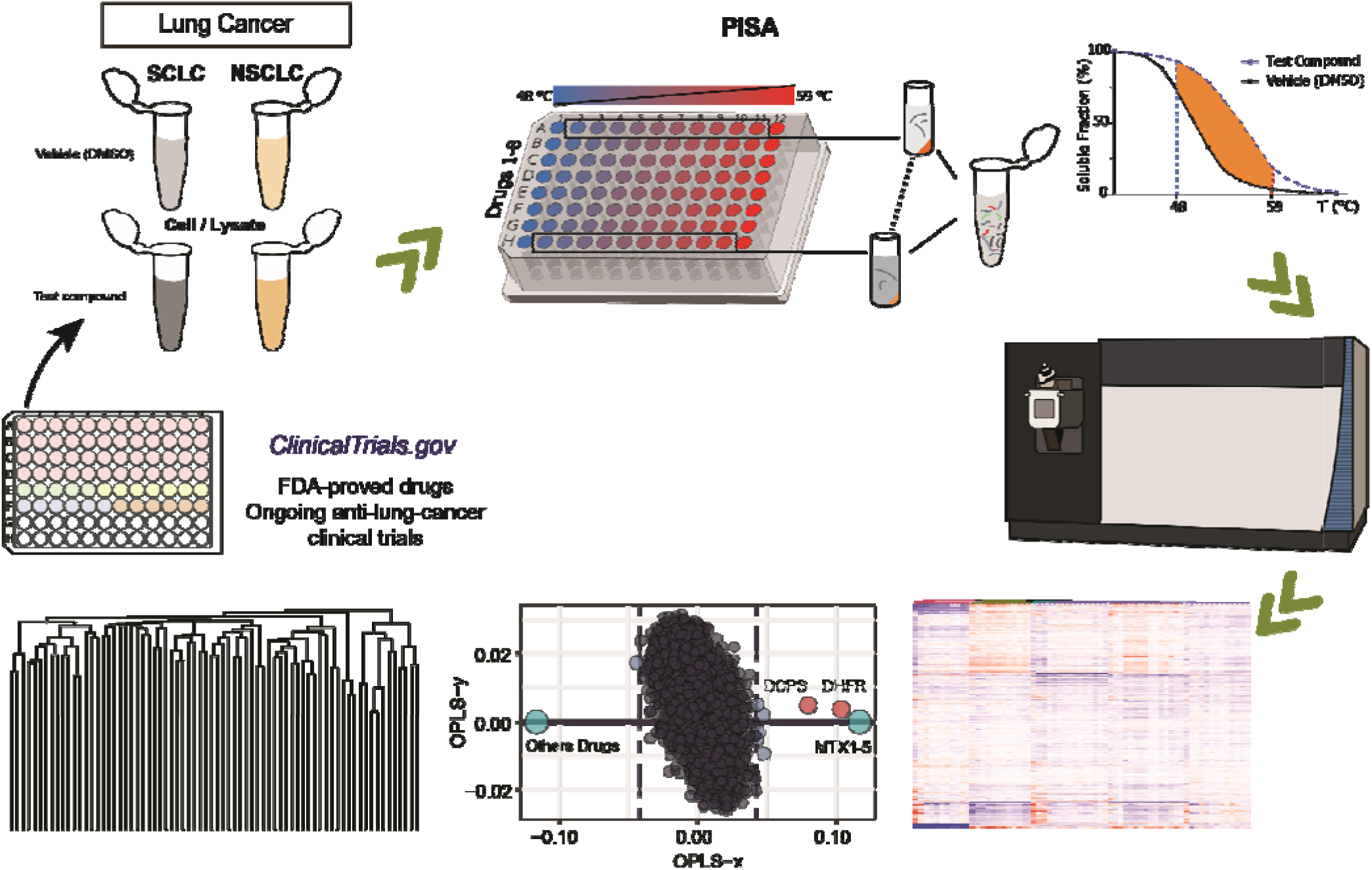
Workflow for PISA-based target identification. The drugs with various known targets and potential mechanisms were chosen from the ongoing chemotherapy-based lung cancer clinical trials. SCLC and NSCLC cell lines were selected as the disease model. PISA analysis was performed both in cell lysate and intact cells. After thermal treatment and ultracentrifugation, proteins were digested, and peptides were labeled with TMT. Samples were then pooled and analyzed by LC-MS/MS. The protein PISA signal shifts compared to vehicle-treated control were visualized on a heat map and processed by OPLS-DA. Upon choosing a proper threshold, a statistical model based on the main OPLS-DA coordinate was used for identification of potential protein targets and hierarchical clustering of the drugs.

To address this, we tested several multivariate analysis methods, including t-distributed stochastic neighbor embedding (tSNE), principal component analysis (PCA), partial least squares discriminant analysis (PLS-DA), and orthogonal partial least squares discriminant analysis (OPLS-DA) (**Supplementary Figure 3**). Among them, OPLS-DA yielded more consistent and biologically meaningful prioritization of known and putative targets across datasets, and was selected as the optimal approach to compare each drug treatment against all other conditions, following the strategy previously applied in expression proteomics^18^. Nevertheless, OPLS-DA of PISA data revealed very few significantly shifting proteins, again in stark contrast with expression proteomics^19^. These puzzling results called for innovative approaches to PISA data processing for reliable identification of the drug targets.

We addressed this unexpected problem as follows. First, from the distribution of proteins’ main OPLS coordinates we estimated for each protein the p-value of being a statistical outlier in that distribution. Then, using the fact that each TMT set had a sample treated with a control drug methotrexate (MTX), a p-value threshold was chosen so that all such samples co-localized most tightly in hierarchical clustering. The outliers in the samples treated with other drugs (typically ≈5% of all quantified proteins) were then considered candidate targets (‘pro-targets’) of a given drug. For validation of the target candidates, we examined the whole dataset: if the same candidate appeared for the same drug in a different type of PISA sample (lysate vs. in-cell, or NSCLC vs SCLC cells), it was considered validated.

## Results and Discussion

### PISA analysis

After the selection of cell lines and drug panel (see methods), cells and cell lysates were treated with the drugs and PISA-analyzed. The number of proteins identified in at least one PISA analysis in A549 cell or lysate samples was 10,632, out of which 9,570 proteins were quantified with at least 2 peptides, excluding potential contaminants. For the H82 cell line, 10,823 proteins and 9,736 proteins were quantified, respectively (**Supplementary Data 1**).

Figure 2 shows the Venn diagram of the overlap between all four datasets encompassing 15 TMT sets. On average, 44% of the proteins detected without missing values were common in all datasets. In A549 cell lysate and cells, 5,063 and 4,626 common proteins were quantified, respectively, while the respective numbers for H82 were 5,650 and 5,756 proteins.

**Figure 2.**
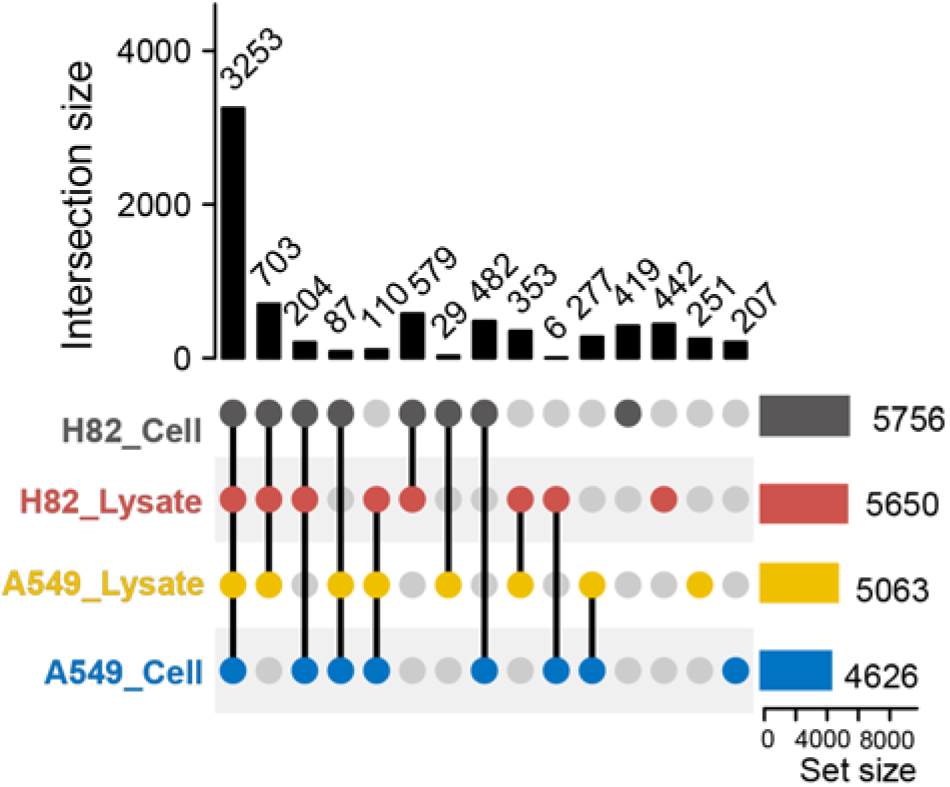
Number of proteins identified and their overlap in the four PISA datasets, A549 cell lysate and cell, and H82 cell lysate and cell. 3253 proteins (expected number is 1872) without missing values were shared across four datasets.

In the first round of analysis, three biological replicates were used to determine the fold-change of the PISA shifts and the respective p-value for each protein under each drug/vehicle condition. The common proteins in a dataset were used for creating a heatmap by hierarchical clustering. As an example, Figure 3A shows a heatmap for the shared proteins from A549 lysate. Heatmaps for the other datasets are shown in **Supplementary** Figure 1**-2.** The first look at the heatmap revealed the already mentioned problem – not all drugs with common known targets co-localized on that heatmap. For instance, selumetinib, trametinib, and binimetinib that all target MEK (MAP2K) were not found in close proximity to each other. At the same time, the PISA shifts of MAP2K1 and MAP2K2 for the three drugs were both strong and statistically significant (**Table 1**), which testifies to the validity of the PISA analysis.

**Figure 3.**
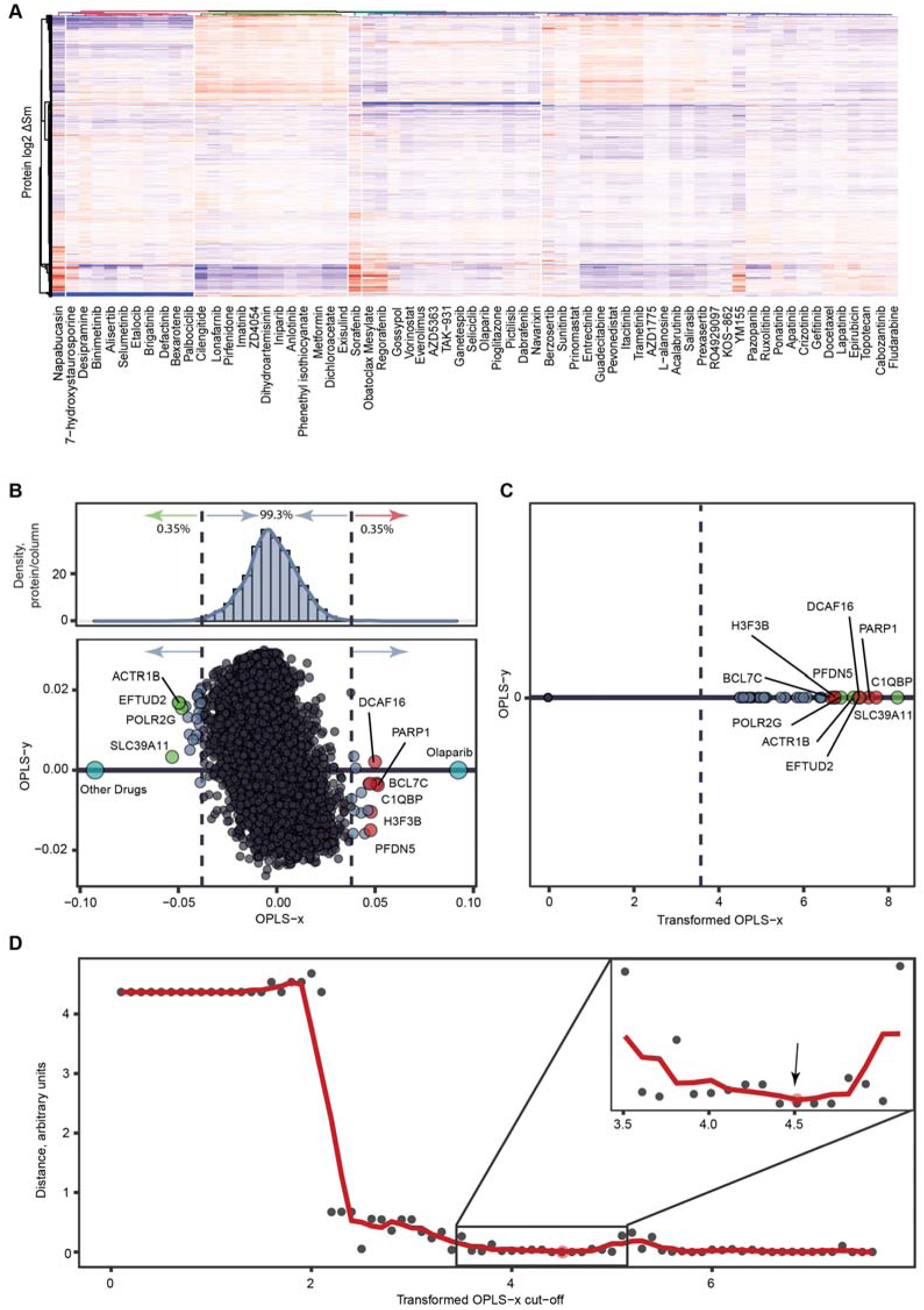
Data processing for drug target identification. **A.** Heatmap based on clustering of log2 transformed PISA fold-changes for 4,626 shared proteins in the A549 lysate dataset. **B.** Lower panel: The loading plot of an OPLS-DA model comparing olaparib to all other drugs in H82 cell dataset. Upper panel: the density distribution of protein’s x-coordinates in the loading plot. Olaparib’s known target PARP1 exhibits the next-highest x-coordinate. **C.** The transformed x-coordinates correspond to -log10(p) of the 39 proteins above the threshold value (dashed vertical line). The top 10 proteins are represented as red/green circles if they became more/less soluble upon olaparib treatment. **D.** Selection of the optimal cut-off threshold. Distance is the average distance between control MTX datasets in hierarchical clustering of all data. Insert shows that the curve reaches its minimum at x=4.5, chosen as optimal cut-off value.

**Table 1.** Fold-changes in PISA and the corresponding p-values of MEK proteins in A549 lysate treated with selumetinib, trametinib or binimetinib vs. DMSO.

|  | Selumetinib |  | Trametinib |  | Binimetinib |  |
| --- | --- | --- | --- | --- | --- | --- |
|  | Fold-change | p-value | Fold-change | p-value | Fold-change | p-value |
| MEK1 (MAP2K1) | 2.3 ± 0.2 | 2.4 × 10<sup>-3</sup> | 2.1 ± 0.1 | 3.5 × 10<sup>-4</sup> | 2.4 ± 0.1 | 1.6 × 10<sup>-4</sup> |
| MEK2 (MAP2K2) | 3.8 ± 0.4 | 4.9 × 10<sup>-3</sup> | 2.4 ± 0.3 | 1.8 × 10<sup>-2</sup> | 4.3 ± 0.5 | 5.7 × 10<sup>-3</sup> |

Examination of the MTX-treated samples (the positive control in each experiment) revealed that they did not cluster together either. The cause of the problem was the specificity of the solubility shift in PISA that affected very few target proteins, while the shifts in the absolute majority of proteins were to a large extent due to statistical fluctuations. Removing this noise and “purifying” the true targets turned out to be a nontrivial but necessary task.

### Optimal p-value threshold

To address this problem, we first applied the OPLS-DA analysis that has demonstrated its utility in the ProTargetMiner expression proteomics database of drug target candidates^18^. Here, the PISA shifts for each drug of interest were contrasted with those for all other drugs and controls (an example for olaparib is shown in Figure 3B, lower panel). The coordinates of each protein along the main axis correspond to a specific solubility increase with drug treatment (positive values) or its decrease (negative values). Compared to the raw PISA shifts, the main OPLS-DA coordinates provide an enhanced specificity, as the shifts common for many drugs obtain a relatively small coordinate value compared to the shifts of similar magnitude that are specific for a given treatment ^18^. We thus expected that the main OPLS-DA coordinate would provide more meaningful clustering of the treatments. There was indeed an improvement, but not sufficient, as the sporadic PISA shifts of unrelated proteins were still posing a problem. It became clear that these unrelated proteins needed to be down-prioritized, so that only proteins passing a certain threshold for statistical significance (outliers) would be used for clustering.

In order to identify the threshold for such statistical outliers, a distribution of the OPLS-DA coordinates was assessed for each model (**Supplementary Data 2**). One example is the OPLS-DA model comparing olaparib vs. all other drugs treated H82 intact cells (Figure 3B). The dispersion of OPLS-DA coordinates was determined, and the p-value for each outlier was calculated using the error function. The latter assumed Gaussian distribution, but the exact shape of the distribution was not critical for the final results. The horizontal axis was then transformed into -log10(p) values (Figure 3C).

Our strategy was to use in further data processing only the PISA shifts for the outliers, zeroing all other PISA shifts that were assumed to be noise. Such an approach required evidence-based determination of the optimal threshold for p-values (dashed vertical line in Figure 3C). For the threshold determination, we used the data on MTX-treated samples that were present in all individual TMT sets (5 such samples in each PISA analysis type). A figure of merit (FoM) function was created corresponding to the average distance in all four types of PISA analysis between the positions of the neighboring MTX samples in hierarchical clustering of all samples of a given type. The tighter the cluster that MTX-treated samples created, the lower FoM was obtained. As Figure 3D shows, the minimal FoM is observed at the value of 4.5. This value was accepted as the optimal -log10(p) threshold for the whole dataset, and the outlying proteins exceeding this threshold, as in Figure 3C, were taken as pro-targets (drug target candidates).

### Drug clustering

With the PISA shifts of all below-threshold proteins set to zero, the hierarchical clustering data became much more meaningful. While the MTX samples clustered tightly as expected (see Figures 4A**, B** as examples), many drugs with similar targets were also found close together. For example, in the A549 lysate dendrogram, the three MEK inhibitors, selumetinib, trametinib and binimetinib, are found next to each other, similar to the potent PTK2 inhibitors brigatinib^20^ and defactinib^21^. With a meaningful clustering achieved, we moved to the identification of the pro-targets and their validation.

**Figure 4.**
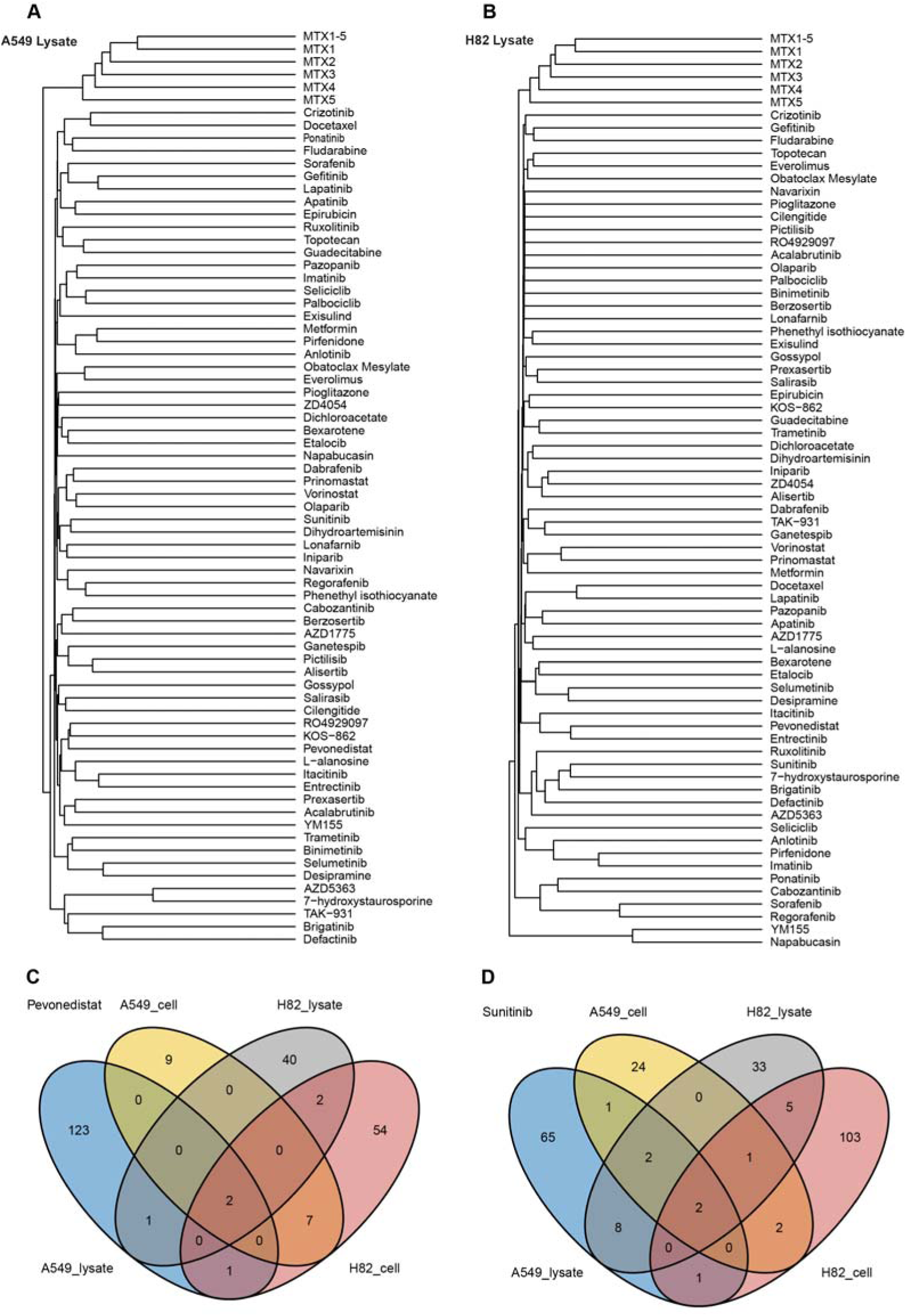
Pro-target-based clustering of the PISA datasets from drug-treated samples together with all MTX-treated controls from the lysates of A549 (**A)** and H82 (**B**) cells upon application of the common optimal threshold of 4.5. Venn diagrams show the overlap of the identified pro-targets of pevonedistat (**C**) and sunitinib (**D**).

### Drug target candidates (pro-targets)

With the optimal threshold of 4.5 for -log10(p), the median number of target candidates for different drugs was 27.5 for A549 lysate and 20 for intact A549 cells, while for H82 cells the median numbers were 24 and 43, respectively. These numbers represent 6.0% of all quantified proteins for A549 lysate and 4.0 % for intact A549 cells, as well as 4.2% for H82 cell lysates and 7.5% for intact cells. For pro-target validation, we applied the following principle: if the same candidate appears in k>1 dataset for the same drug, it is considered validated at the k-th level. All validated pro-targets of 67 drugs are listed in **Table 2**. On average, we found 3 pro-targets per drug with k=2 (overlaps in two datasets), 0.4 pro-targets with k=3 and 0.2 pro-targets with k=4. For control of the false discovery rate (FDR), the ‘candidates’ were selected at random; the numbers of such spurious overlaps were 1.3 proteins with k=2, 4.4×10^-3^ with k=3 and 5.5×10^-6^ with k=4. While these numbers indicate that, strictly speaking, only pro-targets with k≥3 are statistically reliable, we decided to consider also the pro-targets with k=2 to minimize the number of false negatives.

**Table 2.**
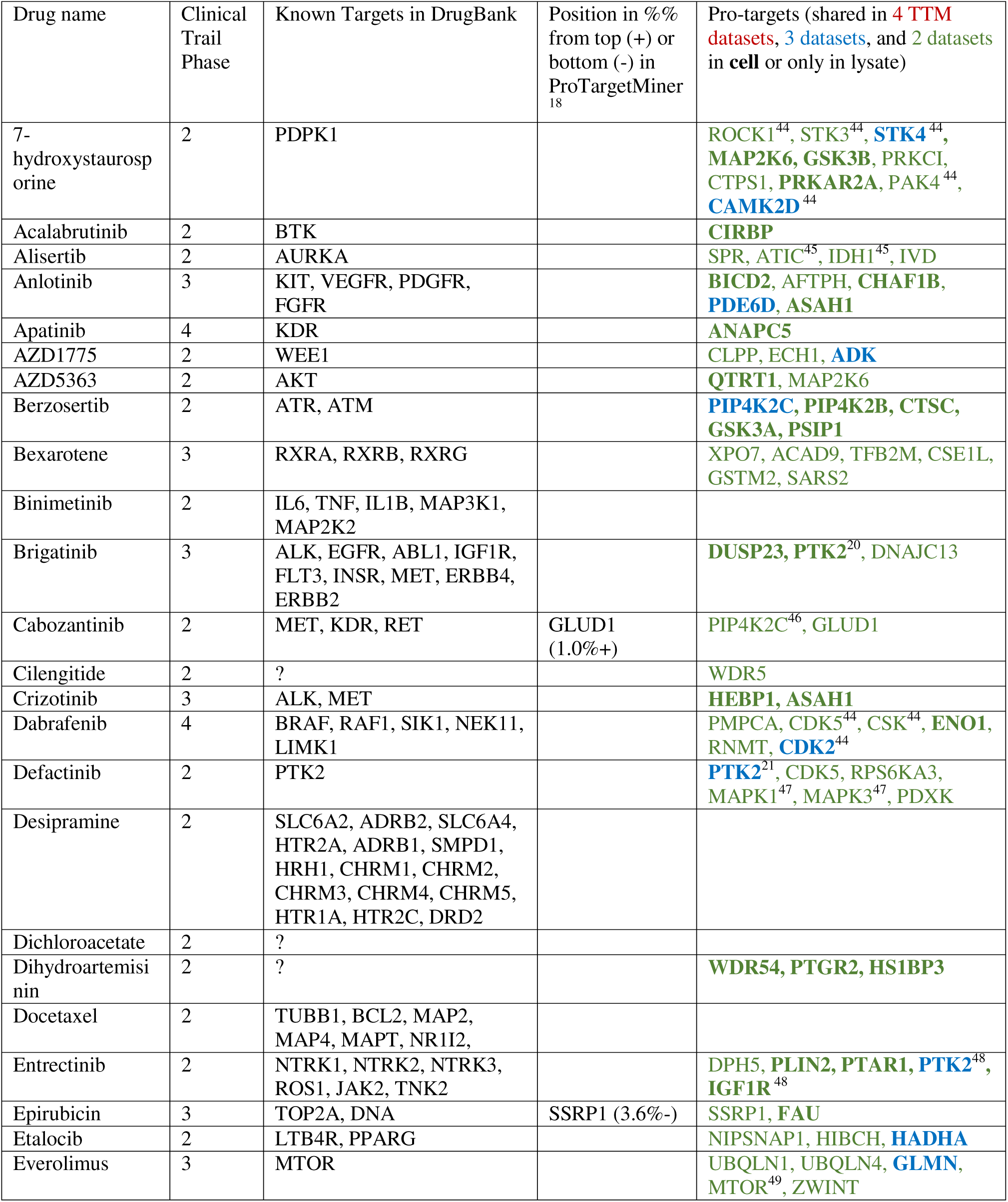
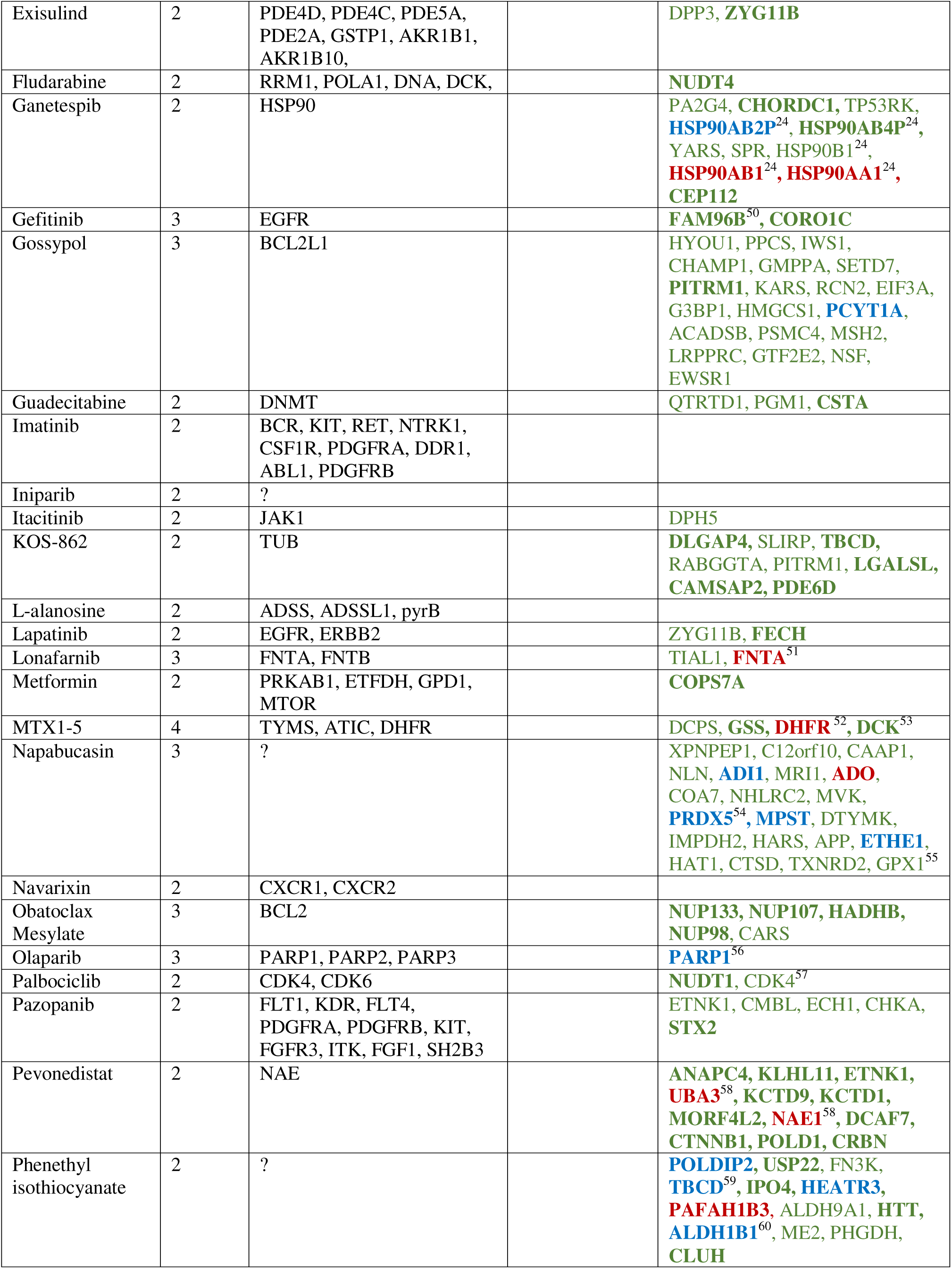
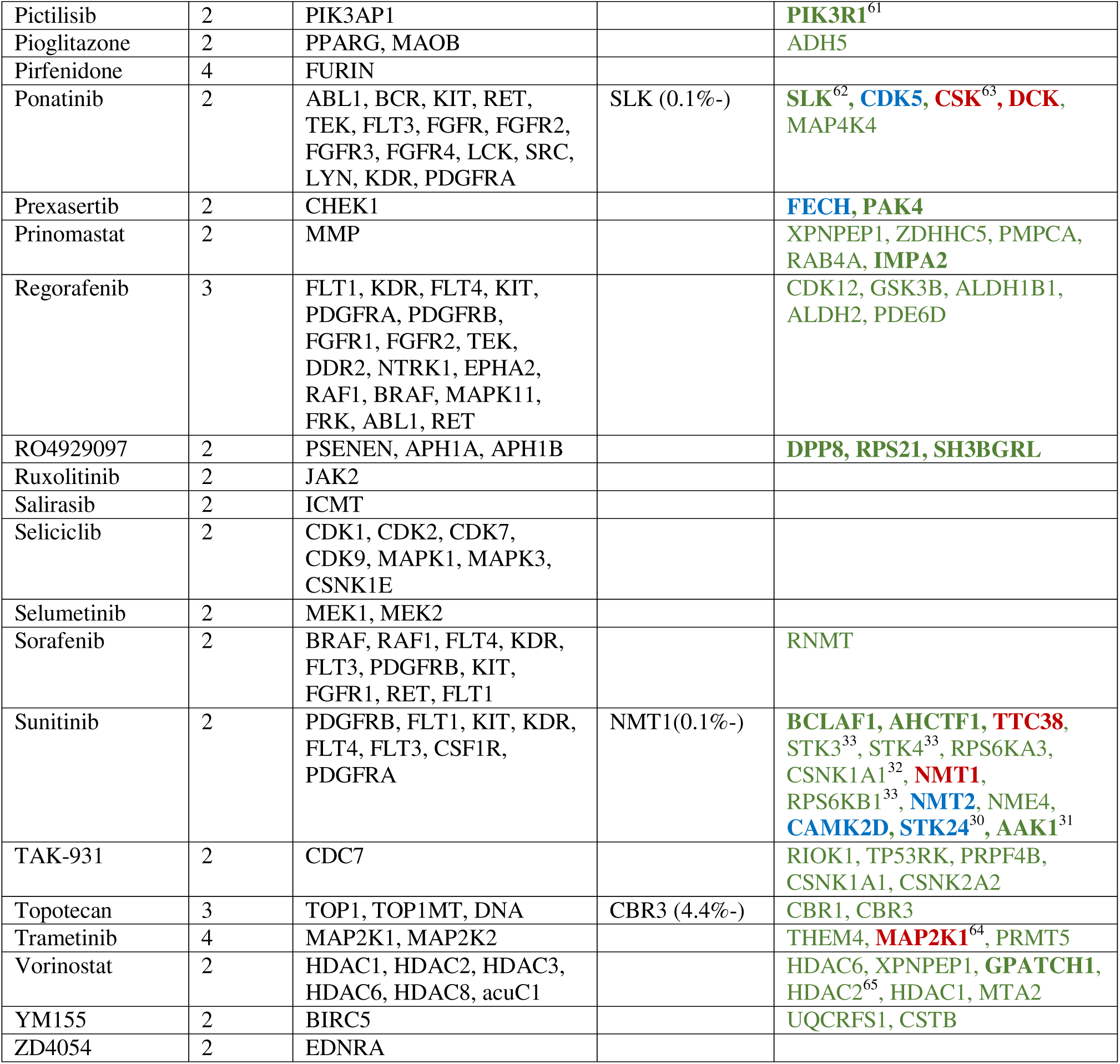
The list of pro-targets validated through our data and ProTargetMiner^18^ for 67 compounds profiled in this study along with clinical phase information and targets of these compounds in DrugBank ^25^.

As an example of pro-target analysis, pevonedistat (also known as MLN4924) treatment in A549 lysate produced 127 target candidates, 18 candidates for intact A549 cells, 45 for H82 lysate and 66 for intact H82 cells (Figure 4C). Of these, 11 pro-targets were found in 2 datasets. Importantly, 2 candidates (NAE1 and UBA3) were found in all 4 datasets, while the anticipated number of randomly shared proteins for k=4 is only 1.7×10^-5^. These two proteins, NAE1 and UBA3 (the latter is also known as NAE2), are two subunits of the known target of that drug - NEDD8-activating enzyme E1 (NAE)^22^. Seven other proteins were shared in two PISA analyses of intact cells. Of these, DCAF7, CRBN and CTNNB1 are found to be co-expressed with NAE^23^.

Another notable example is ganetespib (STA9090), for which 33, 41, 21 and 33 targets candidates were found in the four datasets, respectively. Of these, 9 proteins (expected number – 0.8) were shared in 2 datasets, 1 protein (2.3×10^-3^ proteins expected) in 3 datasets and 2 proteins, HSP90 subunits HSP90AA1 and HSP90AB1, were shared in all datasets (expected number - 2.3×10^-6^). As ganetespib is designed to be an HSP90 inhibitor^24,25^, this result clearly demonstrates the analytical power of the ThermoTargetMiner approach in drug target identification.

These and other target candidates are listed in **Table 2**. The largest number of PISA datasets (20 in total) was obtained for MTX. The DrugBank^25^ lists three MTX targets: TYMS, ATIC, DHFR. Of these, only DHFR is consistently revealed as a pro-target by ThermoTargetMiner. TYMS is an outlier in the MTX-treated intact cells, but not in lysates (Figure 5), which is similar to our previous PISA results^12^. ATIC showed a similar tendency. An explanation for the phenomenon is that, unlike direct binding of MTX to DHFR, MTX needs first to be transformed into MTX polyglutamates (MTXPGs) to exert effects on TYMS and ATIC^26^. Apparently, MTX binding to TYMS and ATIC requires intact cellular environment which provides functional enzymes and substrates. This result confirmed the sufficiently high reliability of the pro-targets identified by PISA for the ThermoTargetMiner database.

**Figure 5.**
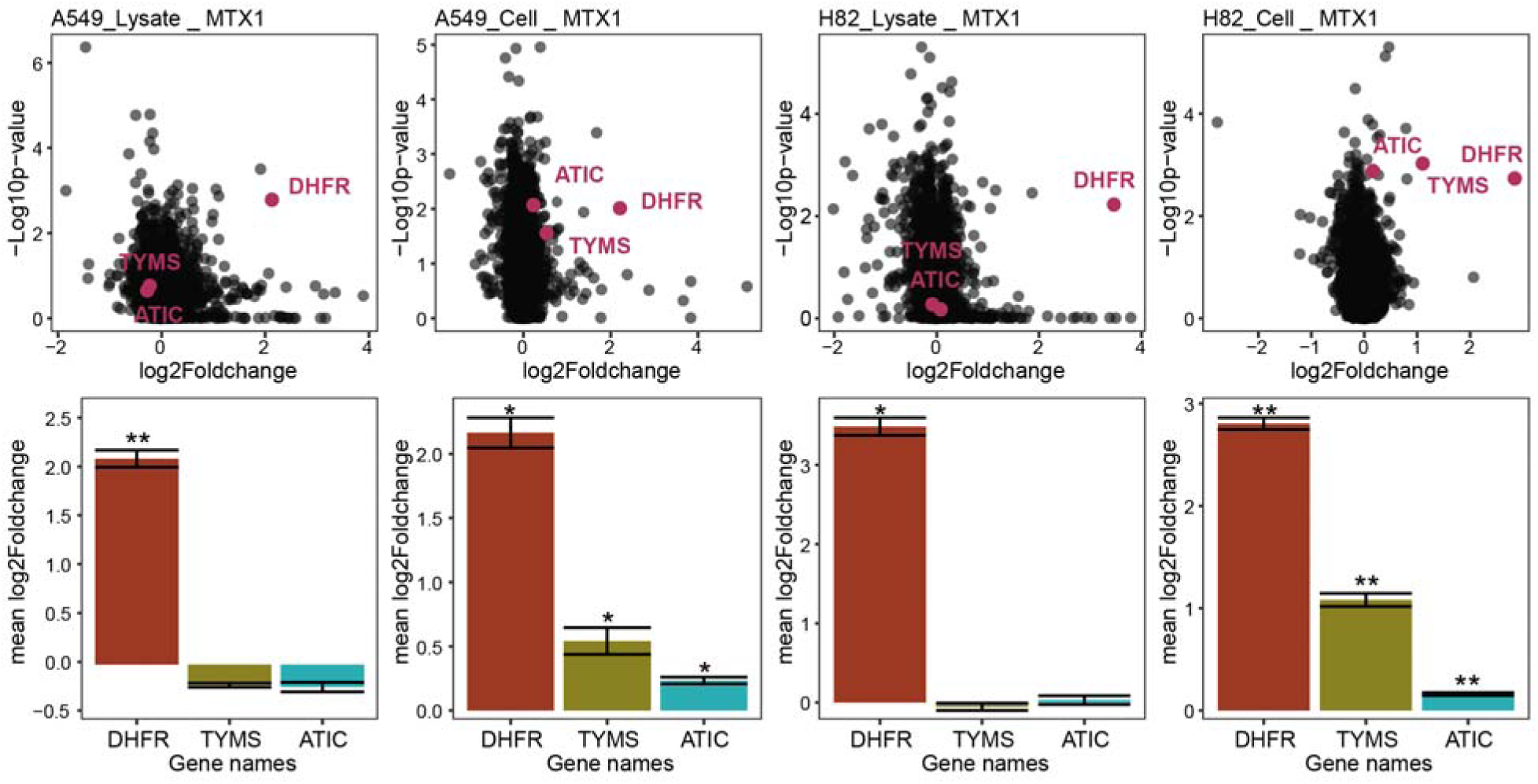
**Fold-changes in PISA and p-values of the three MTX targets listed in Drugbank**^25^**: DHFR, TYMS and ATIC.** * - p < 0.05; ** - p <0.005; *** - p < 0.0005.

### Novel targets

Sunitinib is designed to target receptor tyrosine kinases (RTK), such as vascular endothelial growth factor (VEGF) receptor and platelet-derived growth factor (PDGF)^27^.

However, these transmembrane proteins were not identified in our PISA analysis, likely due to their poor solubility and extraction. Instead, 22 other proteins were shared between at least two datasets (Figure 4D). Among these pro-targets, NMT1 and NMT2, which have been found overexpressed in cancers and thus hinted to be potential anticancer targets^28^, were found in 4 and 3 ThermoTargetMiner datasets, respectively. Previous work^18^ of our group has discovered that sunitinib treatment induces significant NMT1 downregulation, supporting the current PISA finding.

Another protein that exhibited significant PISA shift in all four sunitinib datasets was TTC38 (tetratricopeptide repeat domain 38), which has not been linked to sunitinib binding. Given the very low *a priory* probability of being an outlier with k=4 (only 3.7×10^-5^ such events are expected by pure chance), it is highly likely that TTC38 is a cognate target of sunitinib. To a large extent, the same applies to CAMK2D which was significantly solubilized by sunitinib treatment in three out of four datasets. Of relevance is that the activity of the related protein CaMKII has been significantly elevated following chronic sunitinib treatment, which suggested a mechanism for sunitinib-mediated cardiovascular dysfunction^29^. Furthermore, our results validated the previously reported sunitinib binding to STK24^30^, AAK1^31^, CSNK1A1^32^, RPS6KB1, STK3, and STK4^33^.

Phenethyl isothiocyanate (PEITC) is a natural anti-cancer compound that is present in many cruciferous vegetables. It is believed to suppress cancer progression through diverse mechanisms like cell cycle arrest at the mitotic phase and induction of apoptosis^34^. On the molecular level, PEITC hinders tubulin polymerization and alters tubulin secondary and tertiary structures^35^. However, there is no validated target of PEITC in the Drugbank^25^. In ThermoTargetMiner, PAFAH1B3 (Platelet-activating factor acetylhydrolase IB subunit alpha1), one of the most frequently overexpressed metabolic enzymes in human tumors^36^, was found as the sole pro-target across all four datasets (the expected number of such events is 6.7×10^-6^) (Figure 6A). To confirm the inhibitory effect of PEITC on PAFAH, recombinant PAFAH was incubated with PEITC at a concentration ranging from 0 to 200 µM. The conversion of PAFAH’s substrate 2-thio PAF to 2-(Trimethylammonio)ethyl (2S)-2-(hexadecyloxy)-3-(sulfanyl)propyl phosphate was significantly reduced in a concentration-dependent manner (Figure 6B). In addition, A549 lysate was incubated with PEITC or P11 (a reported inhibitor of

**Figure 6.**
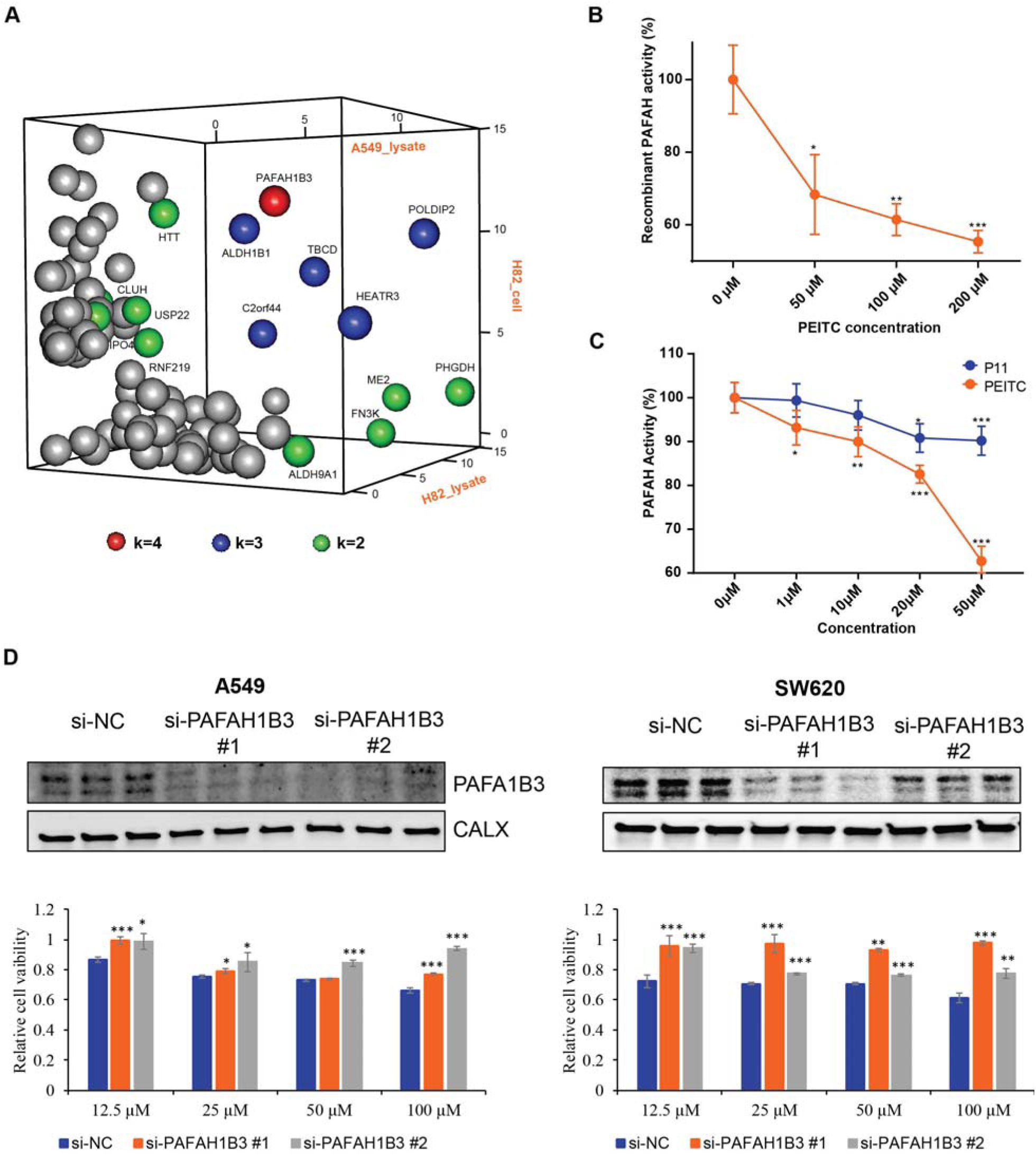
**A.** -log 10(p) of PEITC pro-targets in A549 lysate and H82 cell datasets. The pro-targets are represented by dots colored by their k values. **B.** The activity of recombinant PAFAH across increasing PEITC concentration. **C.** The activity of PAFAH in A549 lysate when incubated with 0-50 µM PEITC or P11. * - p < 0.05; ** - p <0.005; *** - p < 0.0005. **D.** A549 and SW620 cells were transfected with either a negative control siRNA or two different pre-designed siRNAs targeting PAFAH1B3, followed by a cytotoxicity assay under PEITC treatment for 72 hours. For each transfection condition, DMSO-treated cells were included as vehicle controls. The transfection efficiency was measured by immunoblotting. Samples were loaded in triplicates. Calnexin served as a loading control. In the bar chart, cell viability upon PAFAH1B3 with negative control (NC) or siRNAs is expressed as mean ± SD of three parallel samples. * p < 0.05; ** p < 0.01; *** p < 0.001, NS, no significant difference.

PAFAH1B2 and PAFAH1B3^37^), at a concentration ranging from 0 to 50 µM. Upon 30-min incubation, much less products from the PAFAH-induced catalysis was found in PEITC-treated samples, demonstrating that PEITC is more potent than P11 (Figure 6C).

To examine whether PAFAH1B3 contributes to the cellular response to PEITC, we silenced PAFAH1B3 using two siRNAs (si-PAFAH1B3 #1 and si-PAFAH1B3 #2) in A549 cells and in another PAFAH1B3-expressing cell line, SW620. After knockdown, the cells were treated with PEITC for 72 hours. Efficient depletion of PAFAH1B3 was confirmed by immunoblotting (Figure 6D**, upper panel**). Notably, PAFAH1B3 silencing markedly reduced the cytotoxicity of PEITC, resulting in a 20–40% increase in cell viability compared with negative control (NC) siRNA-treated cells (Figure 6D**, lower panel**). These findings confirmed that PAFAH1B3 acts as a functional target mediating the cytotoxic effects of PEITC.

Furthermore, in PEITC data tubulin-specific chaperone D (TBCD), which plays a crucial role in tubulin complex assembly^38^, was an outlier in three datasets (6.3×10^-3^ such events are expected). TBCD is a validated pro-target of one other drug in our database. The drug is KOS-862 (also known as epothilone D or desoxyepothilone B), a tubulin stabilizer known to arrest the cell cycle at the mitotic phase^39^. Interestingly, the effect of KOS-862 on microtubules is opposite to that of PEITC: while PEITC blocks microtubule polymerization^35^, KOS-862 promotes the latter process, facilitating the formation of multipolar spindles^39^. Consistent with that, in PISA analysis these two drugs demonstrate opposite effects on TBCD’s solubility: PEITC treatment increases it, whereas KOS-862 lowers TBCD’s solubility.

### Mechanism of action

Napabucasin (BBI-608) is a novel STAT3 signaling inhibitor that binds to STAT’s hinge pocket and diminishes STAT3 DNA binding affinity^40^. In ThermoTargetMiner, multiple oxidative stress-related proteins, including ADO, ADI1, PRDX5 and ETHE1, showed significant solubility alteration in at least three datasets. Moreover, napabucasin impacted two pivotal regulators of redox homeostasis in humans, thioredoxin and the glutathione system^41^, as both thioredoxin reductase TXNRD2 and glutathione peroxidase GPX1 demonstrated decreased solubility in the two cell lysates. Therefore, we hypothesized that napabucasin acts as an anticancer compound by inducing oxidative stress on cancer cells.

To test this hypothesis, we performed GO enrichment analysis on the PISA data from napabucasin treated A549 cells. The response to oxidation was found to be the most significantly involved biological process (Figure 7A). In agreement with that, it has been reported that napabucasin’s induction of ROS in multiple cell lines^42^ is one of the anti-tumor action mechanisms of this drug. In the same study, napabucasin was found to be a substrate of another oxidoreductase, NAD(P)H dehydrogenase [quinone] 1 (NQO1)^42^. In our A549 lysate and intact cell data, NQO1 was one of the most shifting proteins (Figure 7B **and 7C**). In H82 data, NQO1 was not quantified, possibly because its transcription level in A549 cells is much higher than that in H82 cells (2554 vs. 12 transcripts per million)^43^. In addition to NQO1, multiple key proteins responsible for the maintenance of cellular redox homeostasis, such as PRDX5, PRDX6 and GPX2 were among the top 0.6% shifting proteins in napabucasin-treated A549 intact cells, and in A549 lysate, the p-values of GPX1 and NQO1 ranked as 10^th^ and 11^th^ lowest, respectively.

**Figure 7.**
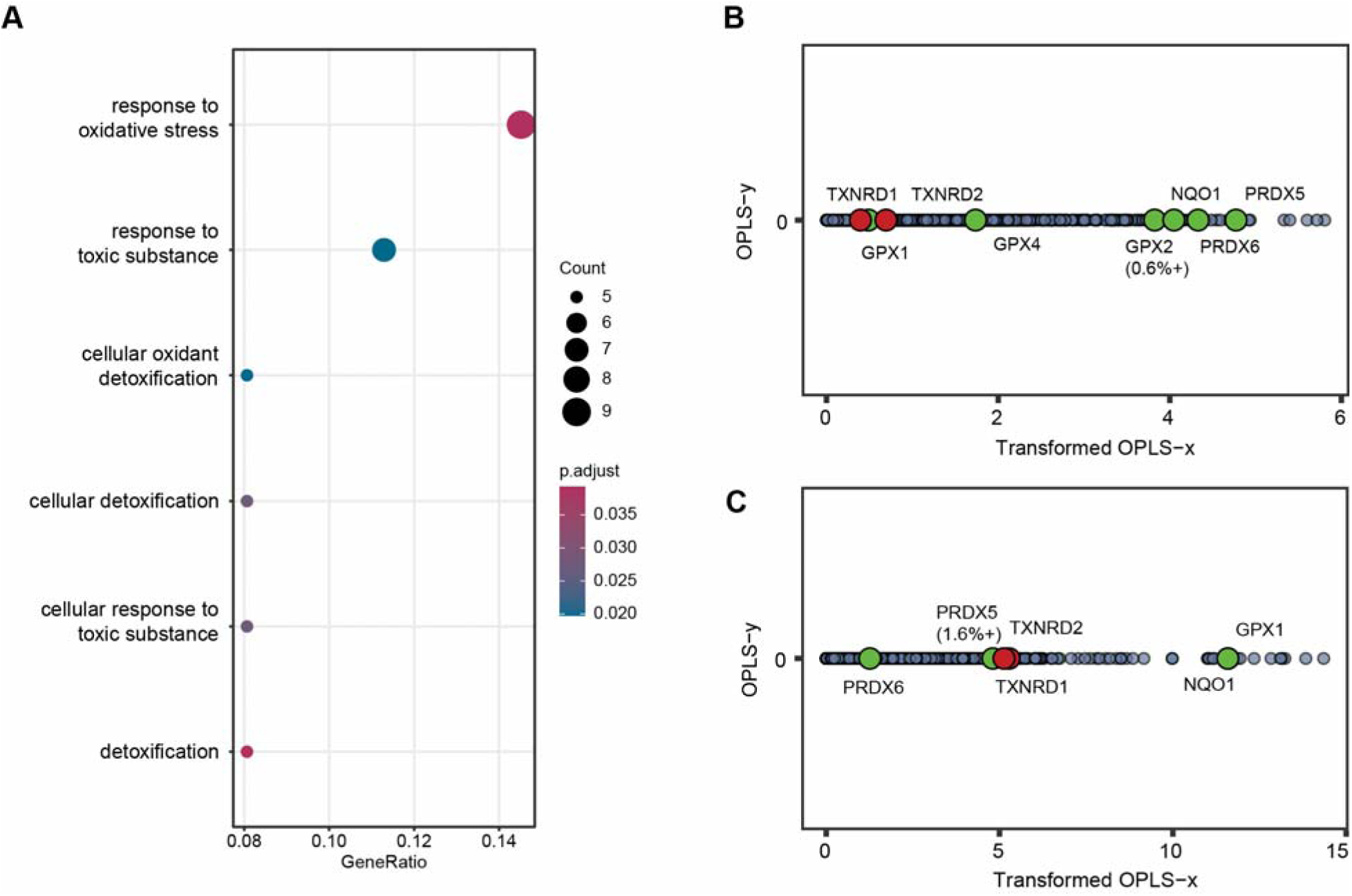
**A**. GO term enrichment in biological pathways of 66 proteins that pass the threshold value of 3 (top 1.4%) in napabucasin treated A549 cells. **B.** Positions on the transformed opls-x scale for A549 cells of the proteins involved in cellular redox homeostasis**. C**. Same for A549 lysate.

### Online Platform for Database Visualization

The ThermoTargetMiner dataset is publicly accessible and can be easily visualized through an interactive R Shiny application. Supported by the SciLifeLab Data Center, the ThermoTargetMiner webpage is hosted on the SciLifeLab Server platform and is available at https://thermotargetminer.serve.scilifelab.se/app/thermotargetminer.

As shown in Figure 8, users can explore pre-built OPLS-DA models for specific drugs in the chosen dataset. By adjusting thresholds, users can define criteria applied for identifying potential drug targets in all four datasets. Proteins that passed the threshold criteria can then be analyzed for pathway enrichment. Common protein targets (pro-targets) of the selected drug are displayed at the bottom of the webpage for easy reference.

**Figure 8.**
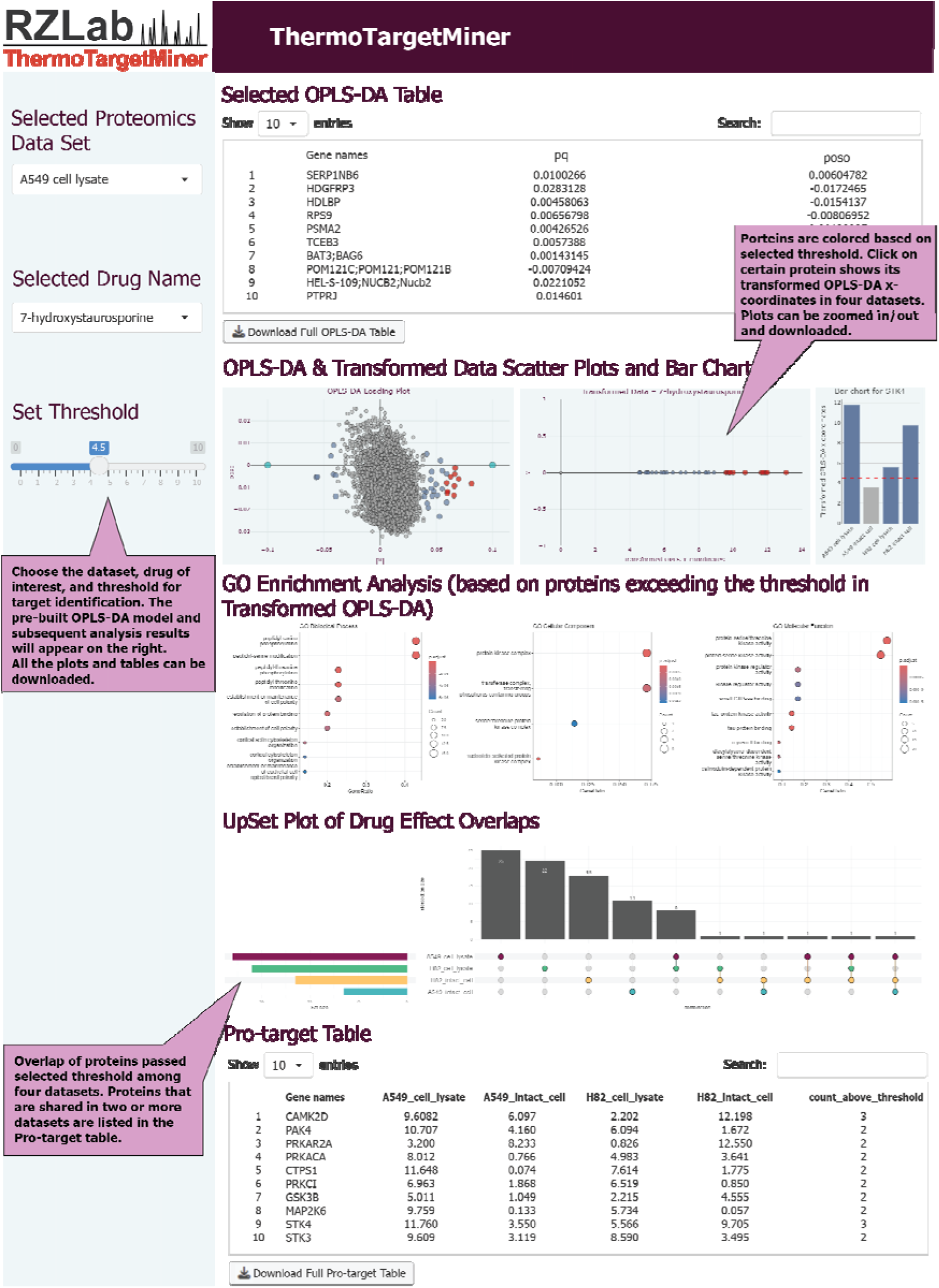
The ThermoTargetMiner R shiny package of target identification and mechanisms.

## Discussion

Identifying drug targets is central to drug discovery and development. Direct or affinity-based approaches are widely used to validate physical drug-protein binding; however, these methods typically require chemical modification of either the drug or the target protein and are often performed outside of a native intracellular environment. In the absence of intact protein complexes, protein conformations may differ substantially from their physiological states, potentially limiting the relevance of observed binding events. In addition, affinity-based methods are inherently biased toward predefined targets and provide limited insight into off-target interactions, which are often critical for understanding drug efficacy and side effects.

Proteomics-based target identification strategies offer a complementary and unbiased alternative, as they do not require engineered probes or prior knowledge of target proteins. Among these approaches, PISA enables detection of drug-protein interactions in a protein-complex-containing environment and is particularly well suited for large-scale applications. By pooling samples across temperature points and leveraging isobaric labeling, PISA achieves an order-of-magnitude increase in throughput compared with traditional thermal proteome profiling (TPP), making it a relatively cost-effective strategy for systematic assessment of on- and off-target drug engagement.

ThermoTargetMiner builds upon these advantages by providing a unified analytical framework for large-scale PISA data. Historically, PISA studies have focused on a limited number of conditions and have relied primarily on protein-wise fold changes and statistical significance for target identification. How well such approaches scale as experimental complexity increases has remained unclear. Previous large-scale PISA efforts have adopted empirical cut-offs based on fold changes and statistical significance, often complemented by extensive biochemical validation. In contrast, our analysis demonstrates that in multi-TMT-set experiments, fold change- and p-value-based methods are particularly susceptible to noise introduced by irrelevant proteins and batch-specific variation. To address this, we introduce normalization combined with OPLS-DA to integrate information across datasets and identify statistically extreme proteins in a multivariate context. Importantly, this strategy enables cross-validation of candidate targets across multiple PISA experiments, reducing reliance on additional validation experiments for every individual protein.

In this study, TMT-based multiplexed quantification with data-dependent acquisition (DDA) was chosen due to its high throughput, mature implementation, and robustness in large-cohort proteomics experiments. DDA workflows with isobaric labeling have well-established software support and enable accurate relative quantification across many conditions within a single run, making them suitable for studies such as ours that compare multiple treatments and biological states. In addition, TMT-based quantification is widely recognized for its strong quantitative precision and typically low coefficients of variation (CVs), reflecting the advantages of multiplexed sample handling and simultaneous MS analysis. However, DDA is known to be limited by its stochastic precursor selection, which preferentially targets the most abundant peptides and can result in incomplete sampling across runs and high missing value rates, particularly for low-abundance peptides or proteins that are inconsistently selected for fragmentation. This bias can also constrain proteome coverage in discovery experiments, as systematic precursor exclusion may overlook informative peptide species that are not among the top selected ions in every cycle.

Over the past few years, data-independent acquisition (DIA)-based approaches have rapidly advanced as an alternative or complement to traditional DDA. Unlike DDA, DIA systematically fragments all precursor ions within defined m/z windows, which improves consistency of peptide detection and reduces missing quantitative values across runs. This feature enhances quantitative reproducibility and can expand proteome coverage, including for low-abundance proteins, when compared to standard DDA workflows. Among emerging strategies, multiplexed DIA (mDIA) has demonstrated promising improvements in proteome depth and quantitative precision, leveraging dimethyl or other labeling schemes combined with reference channels to increase throughput while maintaining comprehensive sampling of peptide ions. Although systematic benchmarking of TMT- and mDIA-based workflows in the context of PISA assays remains to be performed, these developments highlight that DIA-based strategies are becoming feasible for large-scale proteomics and may offer advantages in future applications.

Alternative PISA implementations based on organic solvents or kosmotropic salts have been reported to reveal complementary subsets of drug-protein interactions, potentially including proteins or complexes that are less sensitive to thermal denaturation. However, these approaches often require additional optimization, especially for intact cells, and may not scale as readily to large compound cohorts. In our experience, the thermal approach has a lower tendency for false negative discovery. Similarly, although ultracentrifugation provides a robust separation of soluble and insoluble protein fractions, plate-based filtration has been applied to replace the ultracentrifugation and seems to be a good alternative to more time-consuming ultracentrifugation in large-scale PISA analysis^66^. In this study, ultracentrifugation was used to ensure consistent recovery and minimal bias across the proteome.

ThermoTargetMiner further extends our previous work on ProTargetMiner, in which FITExP was used to infer drug targets based on drug-induced expression changes. While many approved drugs target proteins, not all exert their effects through modulation of protein abundance. For example, kinase inhibitors primarily act by altering phosphorylation states, and small-molecule binding can induce conformational changes without affecting expression levels. PISA therefore serves as a complementary approach to FITExP by capturing solubility changes associated with structural or complex-level alterations. In this study, we establish PISA as a standardized pipeline for large-scale drug screening and provide a curated resource covering 67 anticancer compounds. Beyond identifying known targets, the dataset reveals previously unreported protein associations that may inform side-effect interpretation and suggest opportunities for drug repurposing.

A key component of ThermoTargetMiner is its accessibility as a community resource. To facilitate reuse of the data, we have developed an interactive web interface that enables users to explore OPLS-DA-based target prioritization across compounds and experimental contexts. The web platform allows visualization and download of prioritized targets, while the underlying protein-level fold changes, p-values, and OPLS-DA coordinates are provided in the Supplementary Data. This design allows researchers from diverse fields, including those beyond oncology, to rapidly assess whether proteins of interest exhibit drug-responsive solubility behavior and whether existing compounds may engage these targets, thereby lowering the barrier for hypothesis generation and follow-up studies.

Although ThermoTargetMiner recapitulates several well-established drug-target interactions reported in DrugBank, including PARP1 for olaparib, HSP90 for ganetespib, and the NAE1/UBA3 complex for pevonedistat, not all annotated DrugBank targets are recovered by the PISA-based analysis. This partial overlap reflects the biophysical and technical characteristics of solubility-based proteome profiling, rather than incomplete target engagement. PISA detects drug-induced changes in protein solubility, which depend on factors such as conformational dynamics, post-translational modifications, complex association, protein abundance, subcellular location, proteome coverage and the overall energetics of a given target protein. Consequently, some bona fide drug targets may not exhibit measurable solubility shifts under the applied conditions. This is particularly relevant for membrane proteins, which are underrepresented due to limited solubilization efficiency, lower proteome coverage, and reduced peptide detectability in standard bottom-up mass spectrometry workflows. Similarly, low-abundance proteins may fall below the quantitative detection limit or be masked by stochastic sampling effects, even when functionally engaged by a compound.

In addition, certain targets may bind ligands without inducing appreciable changes in global solubility, especially if binding does not substantially perturb protein folding or higher-order assemblies. Together, these factors contribute to incomplete overlap with DrugBank annotations and underscore that ThermoTargetMiner is not intended to exhaustively reproduce known target lists, but rather to reveal proteins whose physicochemical behavior is measurably altered upon compound treatment in a native or near-native context. This said, ThermoTargetMiner uncovers many new pro-targets and off-targets for these compounds which were not previously covered in databases.

Moreover, some pro-targets may represent indirect or downstream effects of compound treatment, such as altered protein-protein interactions or pathway-level responses, rather than direct physical binding events. While such effects complicate one-to-one comparison with curated target databases, they are nonetheless biologically informative and may be particularly relevant for understanding polypharmacology. Importantly, ThermoTargetMiner distinguishes between solubility shifts observed in intact cells and in cell lysates. Pro-targets detected exclusively in lysates are more likely to reflect direct binding under controlled conditions, whereas those identified only in intact cells may additionally depend on cellular context, metabolism, or protein complex organization.

## Conclusions

This study illustrates how higher-throughput PISA analysis can validate known targets, provide new target information, and help explain the mechanism of drug action. This approach offers a valuable framework for forecasting potential side effects and repurposing drugs for prospective indications. Last but not least, the wealth of target information provided in the ThermoTargetMiner resource holds broader implications beyond lung cancer, and can be extrapolated to various cancer types, to the benefit of a wider oncological community.

## Methods

### Selection of molecules

Clinical trial data for lung cancer were downloaded from https://clinicaltrials.gov/. Only compounds under clinical phase II and above were considered. The complete list of compounds is shown in **Table 1**. Methotrexate was included as a single, well-characterized positive control in each TMT set to monitor assay performance and define an internal reference for statistical optimization. 14 compounds found in ProTargetMiner (crizotinib, docetaxel, ponatinib, sorafenib, sunitinib, gefitinib, lapatinib, pazopanib, ruxolitinib, apatinib, cabozantinib, fludarabine, topotecan and epirubicin)^18^ were selected, along with 53 compounds chosen based on commercial availability and diversity of targets (at least one drug against each known target was included).

### Selection of cell lines

A549 lung adenocarcinoma cells representing NSCLC and NCI-H82 [H82] cells representing SCLC were selected as model systems. A549 is a widely used lung cancer model system that was employed in at least 485,000 studies reported in Google Scholar, including ProTargetMiner^18^. NCI-H82 [H82] was chosen as a classic model of SCLC, because it is adherent and can be grown in the same medium (DMEM) as A549 cells. As this cell line is derived from a metastatic site, it is a good candidate for comparison of the results with A549 cells. None of these cell lines are found in the Register of Misidentified Cell Lines (https://iclac.org/databases/cross-contaminations/).

### Dose and duration of treatment

As compound screening assays for hit discovery are typically run at 1–10 µM^18^, the same concentration of 10 µM was used for all the compounds. The incubation time in the lysate experiments was 30 min, while the cells were treated for 1 h to allow extra time for drug import or diffusion through the cell membrane.

### Proteomics experimental design

In each experiment, two types of controls were used: cells treated with vehicle (DMSO) and with methotrexate (MTX). MTX targets the dihydrofolate reductase (DHFR) protein, which is readily identified in both PISA^12^ and FITExP^19^. The assignment of each TMT channel to each treatment is shown in **Supplementary Table 1.**

### PISA in lysate

PISA experiments were performed using the previously published method^12^. A549 and H82 cells were cultured in 175 cm^2^ flasks, and were then detached, washed twice with PBS, and resuspended in PBS. The cell suspensions were freeze-thawed in liquid nitrogen 5 times, and then centrifuged at 10,000 g for 10 min to remove the cell debris. The protein concentration in the lysate was measured using Pierce BCA assay (Thermo). The cleared lysate was then aliquoted in 3 replicates and treated with the drugs for 30 min at 37°C in 300 μL reaction volume. The 30-minute incubation time was chosen based on literature precedent in PISA and thermal profiling workflows and empirical experience, balancing adequate drug-protein binding equilibration in lysates with the avoidance of prolonged incubation-induced instability. After the reaction, the samples from each replicate were aliquoted into 10 wells in a 96-well plate and heated for 3 min in an Eppendorf gradient thermocycler (Eppendorf; Mastercycler X50s) in the temperature range of 48-59°C. The temperature window of 48–59 °C was selected based on prior thermal profiling studies^12,67^ and preliminary optimization to maximize sensitivity while maintaining adequate proteome coverage. Samples were then cooled for 3 min at room temperature (RT) and afterwards snap frozen and kept on ice. Samples from each replicate were then combined and transferred into polycarbonate thickwall tubes and centrifuged for 20 min at 100,000 g and 4°C.

The soluble protein fraction was transferred to new Eppendorf tubes. Protein concentration was measured in all samples using Pierce BCA Protein Assay Kit (Thermo), the volume corresponding to 25 µg of protein was transferred from each sample to new tubes and urea was added to a final concentration of 4 M. Dithiothreitol (DTT) was added to a final concentration of 10 mM and samples were incubated for 1 h at RT. Subsequently, iodoacetamide (IAA) was added to a final concentration of 50 mM and samples were incubated at RT for 1 h in the dark. The reaction was quenched by adding an additional 10 mM of DTT. No protein precipitation was performed, to avoid losing short semi-tryptic peptides at this stage. Lysyl endopeptidase (LysC; Wako) was added at a 1:75 w/w ratio and samples incubated at RT overnight. Samples were diluted with 20 mM EPPS to the final urea concentration of 1 M, and trypsin was added at a 1:75 w/w ratio, followed by incubation for 6 h at RT. Acetonitrile (ACN) was added to a final concentration of 20% and TMT reagents were added 4x by weight (200 μg) to each sample, followed by incubation for 2 h at RT. The reaction was quenched by the addition of 0.5% hydroxylamine. Samples within each replicate were combined, acidified by TFA, cleaned using Sep-Pak cartridges (Waters) and dried using DNA 120 SpeedVac Concentrator (Thermo). The pooled samples from each of the five TMT experiments were resuspended in 20 mM ammonium hydroxide and fractionated by offline high-pH reverse-phase chromatography. The fractionation system was a Dionex Ultimate 3000 2DLC system (Thermo Scientific) carrying an XBrigde BEH C18 2.1×150 mm column (Waters; Cat#186003023), and the fractionation was done over a 48 min gradient of 1-63% B (B=20 mM ammonium hydroxide in acetonitrile) in three steps (1-23.5% B in 42 min, 23.5-54% B in 4 min and then 54-63% B in 2 min) at 200 µL min-1 flow. Fractions were then concatenated into 12 samples in sequential order (e.g., fractions 1, 13, 25, …, and 85 were combined).

### PISA in cells

Cells were cultured in 6-well plates to a density of 250,000 cells per plate. A day later, cells were treated with the drugs for 1 h. The cells were then washed with PBS, scraped off and resuspended in PBS. The cells were then aliquoted into 10 in PCR plates and heated like above. The cells were then snap-frozen and kept on ice. The samples from each replicate were then pooled and 0.4% final concentration of NP40 was added. The rest of the protocol was identical to PISA in lysate.

### LC-MS/MS analysis and data acquisition

Orbitrap Fusion and Lumos mass spectrometers were used online with an Ultimate 3000 RSLC nanoUPLC system (Thermo Scientific). Sample fractions were dried and resuspended in Buffer A (0.1% FA and 2% acetonitrile in water) to a theoretical peptide concentration of 0.3 µg/µL. Resuspended peptides were loaded onto a Acclaim PepMap 100 C18 HPLC column (75 µm internal diameter, 3 µm beads, 100 Å pore size, Thermo, Cat#164535) for 5 min at a flow rate of 4 μL/min. Peptides were transferred through an EASY-Spray column (75 µm internal diameter, 2 µm beads, 100 Å pore; Cat#ES903) connected to the Easy-Spray source (Thermo; Cat#ES082). Subsequently, the peptides were eluted with a buffer B (0.1% FA and 2% water in acetonitrile) gradient at a flow rate of 300 nL min^−1^. The elution gradient was from 4% B to 28% B for 150 min, to 34% B for 15 min, increasing to 95% B in the next 3 min and staying at 95% for 4 min. Mass spectra were acquired with an Orbitrap Fusion Tribrid mass spectrometer (Thermo; Cat#IQLAAEGAAPFADBMBCX) in the data-dependent mode with MS1 analysis at 120,000, and MS2 at 50,000 resolution, in the m/z range from 400 to 1600. Peptide fragmentation was performed via higher-energy collision dissociation at 35% normalized collision energy.

### Protein identification and data analysis

Raw LC-MS/MS data were processed for protein identification and quantification using MaxQuant software (2.5.0.0) with the UniProt human proteome database (UP000005640_9606 and UP000005640_9606_additional). No more than two missed cleavages were allowed. And the results were filtered to a 1% false discovery rate. Data post-processing was performed in R and OPLS-DA models were built using SIMCA 17 (Sartorius).

### Statistics

Only proteins that were identified with two or more unique peptides and without potential contaminations were included in the final dataset. To calculate the fold-changes of the PISA signals, abundances of TMT reporters (peptide abundances) were first normalized to the total abundance in each TMT channel, followed by the protein abundances being calculated as the average of all normalized peptide abundances. Thereafter the protein abundances were normalized to those in the DMSO-treated samples (occupying the TMT126 channel in each TMT set, see Supplementary Table 1). The fold-changes were then calculated as the ratios of the protein abundances in treated samples vs those of the controls. Batch effects among three biological replicates were removed using Limma package^68^. For each protein, the median fold-change was used for further analysis, and the p-values were calculated by the two-sided Student’s t-test on the normalized abundances in treated samples vs those in controls. In OPLS-DA analysis, protein coordinates were normalized first to the coordinates of the drug, and then to the standard deviation (SD) of the distribution of proteins’ x-coordinates. The OPLS-DA-derived p-values were the GAUSS error function calculated for each protein based on its SD-normalized coordinate.

### LC-MS/MS Data availability

The LC-MS/MS raw data files and extracted peptides and protein abundances are deposited in the PRIDE repository of the ProteomeXchange Consortium^69^ under the dataset identifier PXD054158 with no restrictions. The source data of protein fold changes are provided as a Source Data file. All other data are available from the corresponding authors on request.

### PAFAH activity assay

We used the PAF Acetylhydrolase Assay Kit (No. 760901, Cayman) for PAFAH activity investigation. A549 cells were cultured in T75 flasks and the lysate preparation was the same as PISA lysate preparation except the cells were harvested using a cell scraper. The cells were washed by PBS once and harvested, then were freeze-thawed for 5 cycles. The lysate was then centrifuged at 10,000 g for 10 minutes. The supernatant is transferred to a new tube. The lysate was incubated with drugs or control (DMSO) at the final concentrations of 1, 10, 20 and 50 µM for 30 minutes at 37 °C. For recombinant PAF-AH + PEITC wells, 24 µL recombinant PAF-AH was mixed with 156 µL PBS and the mixture was subsequently divided into three aliquots. PEITC was added to the final concentration of 50, 100 and 200 µM ahead of 30 minutes of incubation at 37 °C. The buffers were prepared, and the reaction was added to a 96-well plate following the kit’s manual. The absorbance data of the two parts - lysate-containing wells and recombinant PAF-AH wells were analyzed separately. The absorbance of drug-treated A549 lysate wells was first normalized to background lysate control and then normalized to the untreated lysate control. For the recombinant PAF-AH incubated with PEITC wells, their absorbance minus absorbance in no-enzyme control wells, then was normalized to absorbance in recombinant PAF-AH control wells.

### siRNA Transfection and Cell Viability Assay

A549 cells were transfected with a negative control siRNA or three different pre-designed siRNAs targeting PAFAH1B3 (MedChemExpress, HY-RS09969) for 24 hours by using Lipofectamine™ 2000 (Thermo Fisher Scientific) according to the manufacturer’s instructions. Transfection efficiency was verified by immunoblotting using a PAFAH1B3 (ProteinTech, 20564-1-AP) antibody, with Calnexin (Enzo Life Sciences, ADI-SPA-860-F) serving as a loading control. Cells were then seeded into 96-well plates at a density of 3,000 cells per well and allowed to adhere overnight. The following day, cells were treated with either DMSO (control) or PEITC for 60 hours. After treatment, Cell Titer-Blue reagent (Promega) was added for 4 hours, and absorbance was subsequently measured using a plate reader.

## Acknowledgements

This work was supported by Cancerfonden (grants 19 0558 Pj and 22 1967 Pj to RAZ). AAS was supported by grants from the Swedish Research Council (2023-02692) and Cancerfonden (24 3595 Pj). We would like to acknowledge Ákos Végvári and Xuepei Zhang for their assistance in LC-MS/MS analysis and Marie Ståhlberg and Carina Palmberg for their general assistance in lab work. We also appreciate Karolinska Institutet for open access funding. RAZ also acknowledges support from The Ministry of Science and Higher Education of RF (agreement № 075-15-2020-899), as well as RUDN project № 033322-2-000.

## Author contributions

The concept, resources, experimental design, and methodology by R.A.Z., protocols and training by A.A.S.; sample preparation by H.L., A.A.S., B.S., A.V., and B.N.; LC-MS/MS analysis by H.L. and Z.M.; siRNA and WB analysis by M.V.; data analysis and visualization by H.L. and H.G.; writing - original draft by H.L. and R.A.Z., with editing by A.A.S. and reviewing by all other co-authors.

