## Supplementary Material for "ThermoTargetMiner as a proteome integral solubility alteration target database for prospective drugs against lung cancer"

**Lyu et al.**

**Supplementary Table 1.** Assignment of the TMT channels in the experiments 1-5 for each of the cell lines (A549 and NCI-H82) and analysis type (in intact cells and lysate).

|  | **Experiment 1** | **Experiment 2** | **Experiment 3** | **Experiment 4** | **Experiment 5** |
| --- | --- | --- | --- | --- | --- |
| **TMT126** | DMSO | DMSO | DMSO | DMSO | DMSO |
| **TMT127N** | Methotrexate | Methotrexate | Methotrexate | Methotrexate | Methotrexate |
| **TMT127C** | Crizotinib | AZD5363 | RO4929097 | Sunitinib | Brigatinib |
| **TMT128N** | Docetaxel | Obatoclax Mesylate | KOS-862 | Prinomastat | Defactinib |
| **TMT128C** | Ponatinib | Gossypol | AZD1775 | Phenethyl isothiocyanate | Bexarotene |
| **TMT129N** | Sorafenib | Dabrafenib | L-alanosine | Metformin | Selumetinib |
| **TMT129C** | Gefitinib | TAK-931 | Prexasertib | Dichloroacetate | Napabucasin |
| **TMT130N** | Lapatinib | Seliciclib | Guadecitabine | Anlotinib | Desipramine |
| **TMT130C** | Pazopanib | Navarixin | Itacitinib | Lonafarnib | 7-hydroxystaurosporine |
| **TMT131N** | Ruxolitinib | Regorafenib | Trametinib | Pirfenidone | Etalocib |
| **TMT131C** | Apatinib | Vorinostat | Pevonedistat | Exisulind | Binimetinib |
| **TMT132N** | Cabozantinib | Ganetespib | Entrectinib | Cilengitide | Alisertib |
| **TMT132C** | Fludarabine | Everolimus | YM155 | Dihydroartemisinin | Palbociclib |
| **TMT133N** | Topotecan | Olaparib | Berzosertib | Iniparib | Empty |
| **TMT133C** | Epirubicin | Pictilisib | Acalabrutinib | Imatinib | Empty |
| **TMT134** | SDS | Pioglitazone | Salirasib | ZD4054 | Empty |

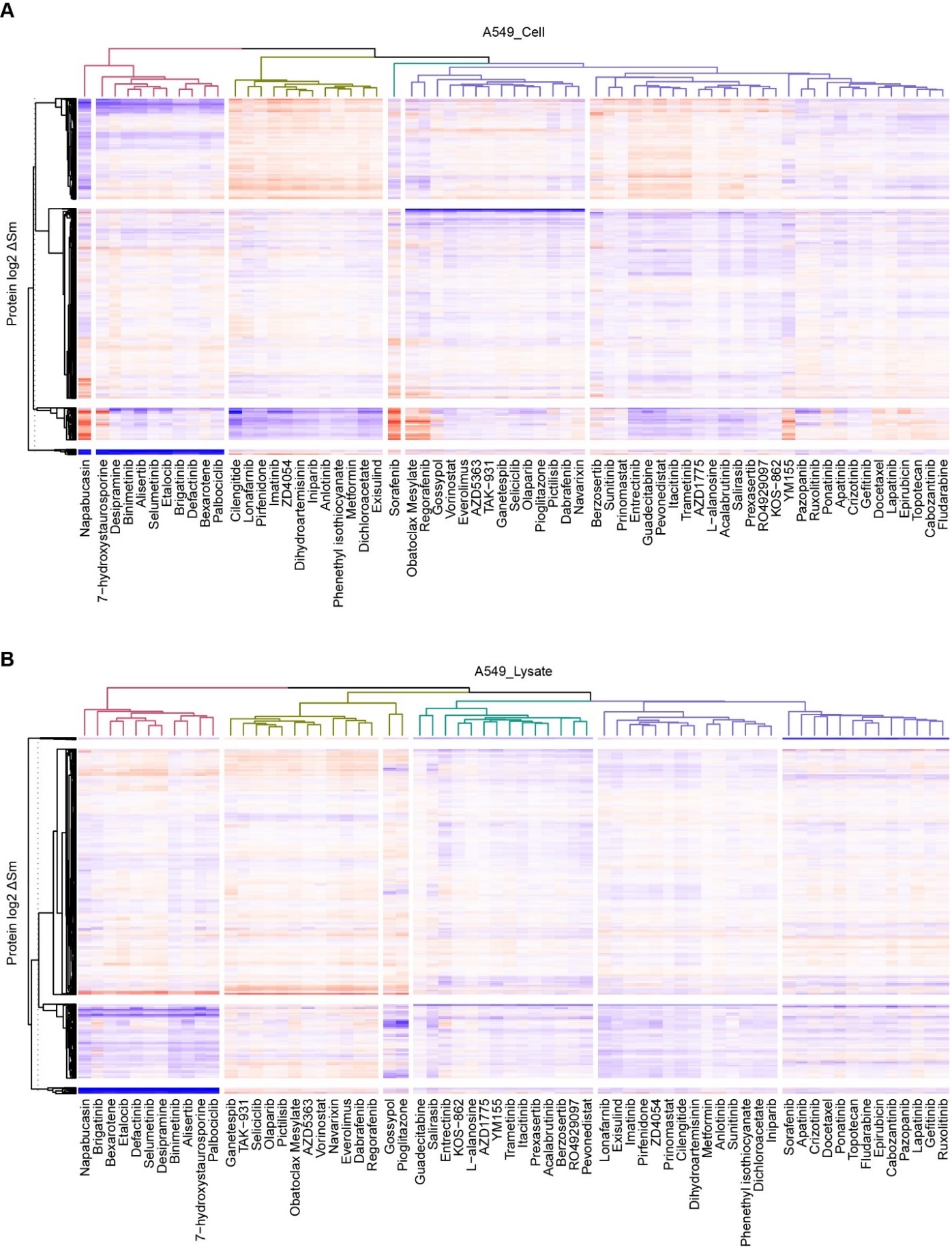

**Supplementary Figure 1.** Heatmap based on clustering of log2 transformed PISA fold changes for proteins in the A549 lysate dataset (**A**) and A549 intact cell dataset (**B**).

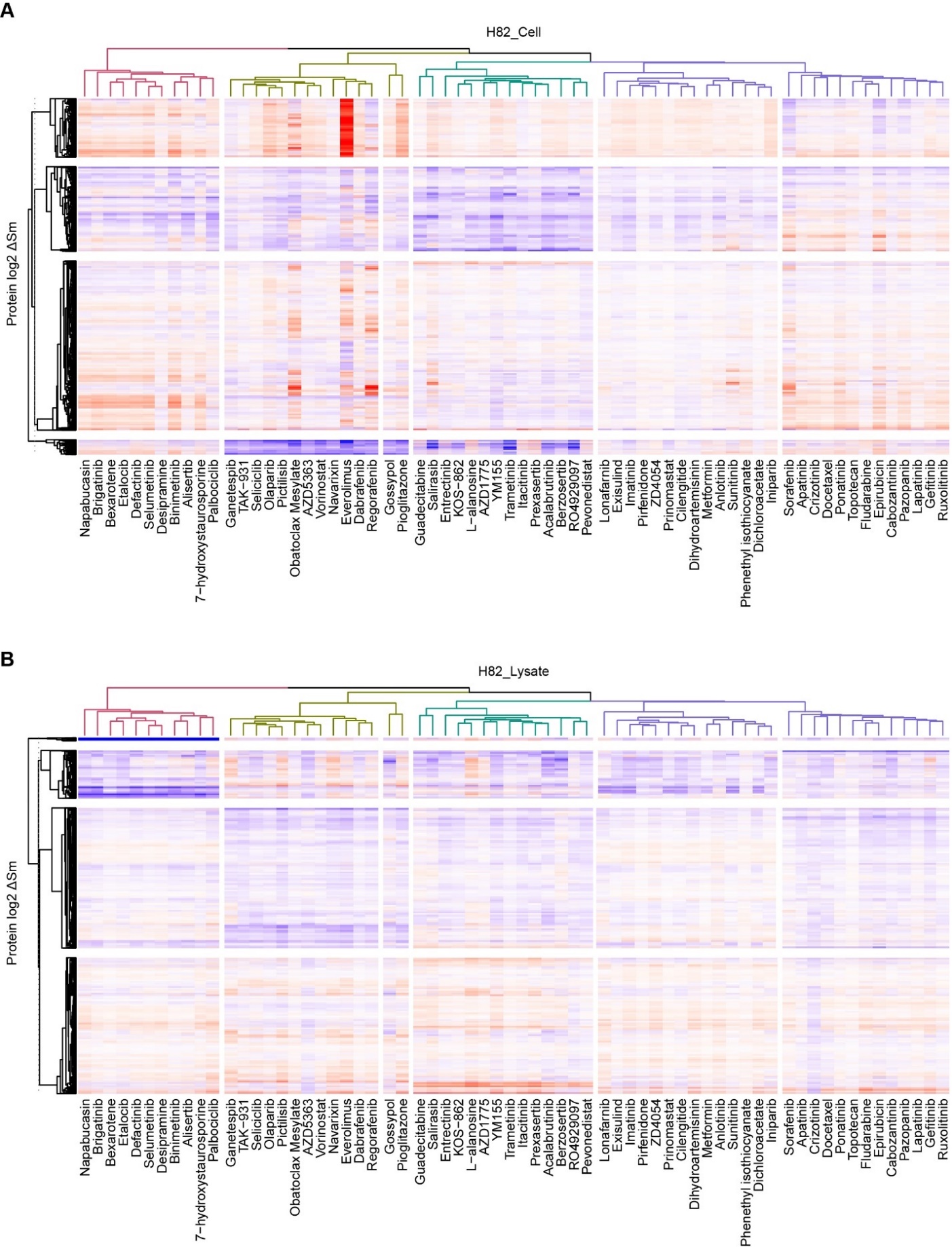

**Supplementary Figure 2.** Heatmap based on clustering of log2 transformed PISA fold changes for proteins in the H82 lysate dataset (**A**) and H82 intact cell dataset (**B**).

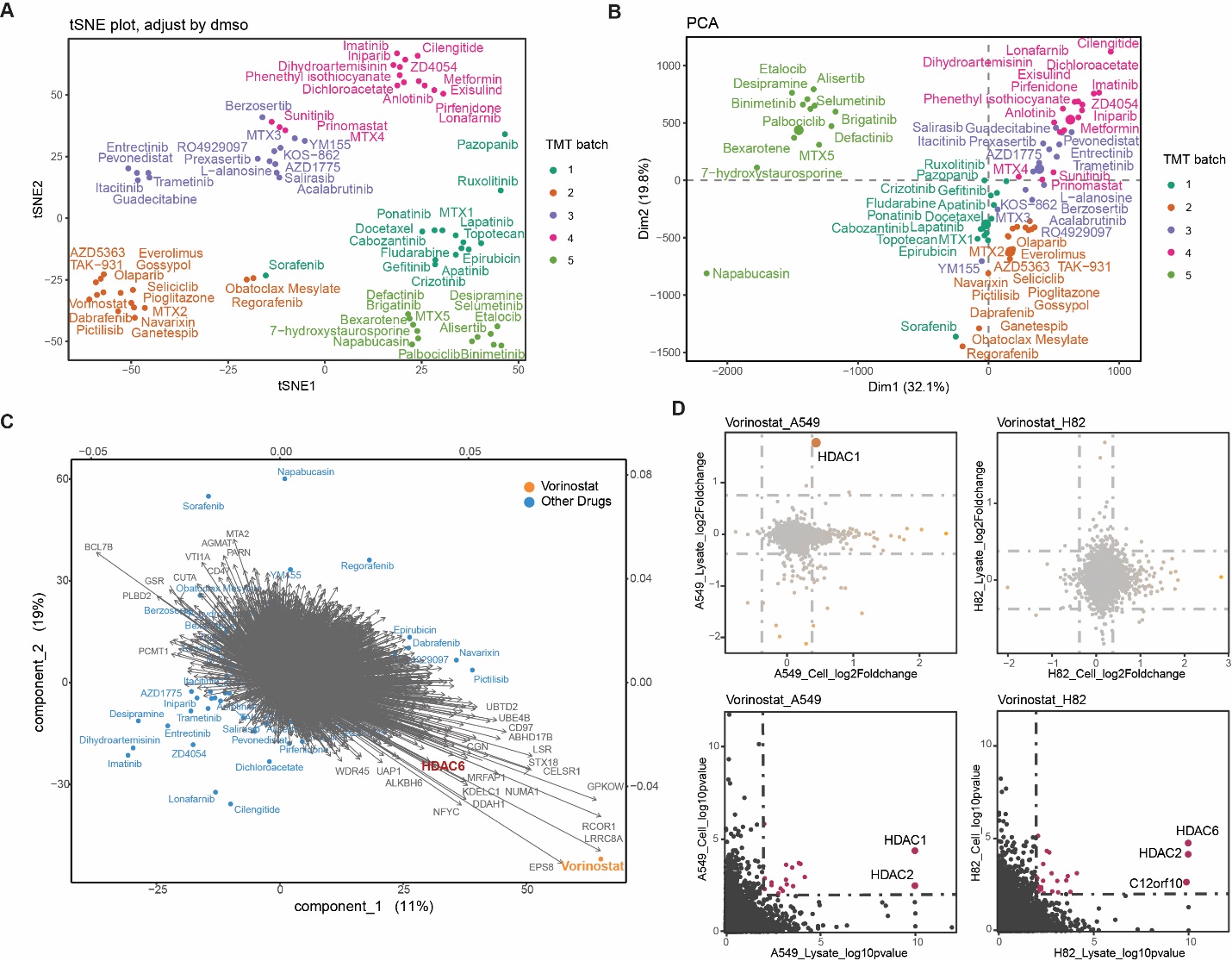

**Supplementary Figure 3.** **Comparison of statistical analysis approaches for ThermoTargetMiner data.** **(A-B) Unsupervised clustering reveals dominant batch effects.** **(A)** t-SNE analysis of A549 PISA data. Despite normalizing drug-treated samples against batch-matched DMSO controls (see Methods), samples cluster primarily by TMT experimental batch (indicated by color) rather than drug mechanism. **(B)** Principal Component Analysis (PCA) of sum- and median-normalized data. Similar to t-SNE, the variance is driven by batch effects ("Set") rather than biological differences, obscuring specific drug-target signals. **(C) Limitations of one-vs-all PLS-DA.** A biplot visualizing a PLS-DA model comparing one specific drug (e.g., vorinostat) against all other drugs. While this approach can identify true targets (e.g., HDACs), it is suboptimal due to projection biases requiring manual adjustment. **(D) OPLS-DA improves target identification sensitivity.** Comparison of target detection using standard thresholding versus OPLS-DA. **Upper panels:** Scatter plots using standard filtering criteria (Fold Change > 1.3 and p-value < 0.05). This method shows low sensitivity; for example, only HDAC1 is identified in A549 lysate/cell data, and no targets passed thresholds in H82 data. **Lower panels:** Application of OPLS-DA scoring and coordinates transformation to the same dataset. This method significantly enhances signal-to-noise separation, successfully identifying known targets (HDAC1, HDAC2, and HDAC6) as top hits.
